# Sequential Activation and Local Unfolding Control Poly(A)-Binding Protein Condensation

**DOI:** 10.1101/2022.09.21.508844

**Authors:** Ruofan Chen, Darren Kahan, Julia Shangguan, Joseph R. Sachleben, Joshua A. Riback, D. Allan Drummond, Tobin R. Sosnick

**Author notes:** Department of Molecular and Cellular Biology, Baylor College of Medicine; Houston, TX, USA.

## Abstract

Eukaryotic cells form biomolecular condensates to sense and adapt to their environment^1,2^. Poly(A)-binding protein (Pab1), a canonical stress granule marker^3,4^, condenses upon heat shock or starvation, promoting adaptation^5^. The molecular basis of condensation has remained elusive due to a dearth of techniques to probe structure directly in condensates. Here we apply hydrogen-deuterium exchange/mass spectrometry (HDX-MS) to investigate the molecular mechanism of Pab1’s condensation. We find that Pab1’s four RNA recognition motifs (RRMs) undergo different levels of partial unfolding upon condensation, and the changes are similar for thermal and pH stresses. Although structural heterogeneity is observed, the ability of MS to describe individual subpopulations allows us to identify which regions become partially unfolded and contribute to the condensate’s interaction network. Our data yield a clear molecular picture of Pab1’s stress-triggered condensation, which we term *sequential activation*, wherein each RRM becomes activated at a temperature where it partially unfolds and associates with other likewise activated RRMs to form the condensate. This model thus implies that sequential activation is dictated by the underlying free energy surface, an effect we refer to as *thermodynamic specificity*. Our study represents a methodological advance for elucidating the interactions that drive biomolecular condensation that we anticipate will be widely applicable. Furthermore, our findings demonstrate how condensation can use thermodynamic specificity to perform an acute response to multiple, stresses, a potentially general mechanism for stress-responsive proteins.

## Main Text

Cells form clusters of RNA and RNA-binding proteins in response to stresses^3,4,6,7^. These clusters, broadly termed stress granules, fall into the category of biomolecular condensates. Studies investigating the molecular basis of condensation typically focus on low complexity regions (LCRs) and intrinsically disordered regions (IDRs) that mediate weak multivalent interactions to drive condensation^8–11^. Yet in important physiological cases, such as for the core stress granule marker poly(A)-binding protein (Pab1 in yeast), condensation is mediated by its folded domains^5^.

Pab1 is consistently recruited to stress granules across a range of stresses^3,4^ including heat shock, starvation, oxidative stress, and osmotic stress. In non-thermal stresses, stress-induced intracellular acidification acts as a second messenger to trigger condensation^5,11–13^. Accordingly, upon heat shock or starvation-induced acidification, Pab1 autonomously condenses even before stress granule formation^5^. The resulting condensates are irreversible by cooling (Extended Data Fig. 1a) but can be dispersed by stress-induced molecular chaperones orders of magnitude faster than misfolded proteins^14^.

Mimicking its *in vivo* behavior, purified Pab1 condenses *in vitro* above a temperature T_cond_=39 °C (at pH 6.4) or below pH 5.4 (at 30 °C), both of which overlap with *in vivo* stress conditions, and smoothly integrates these two signals into a single phase boundary (Fig. 2b)^5^. Notably, the condensation of Pab1 is adaptive with suppression of its condensation reducing cellular fitness during stress; potentially by altering the translations of housekeeping and stress-responsive transcripts^15,16^. However, crucial questions remain: What is the structural basis of Pab1’s stress-triggered condensation? How does Pab1 integrate thermal and pH signals into a single phase boundary?

**Fig. 1.**
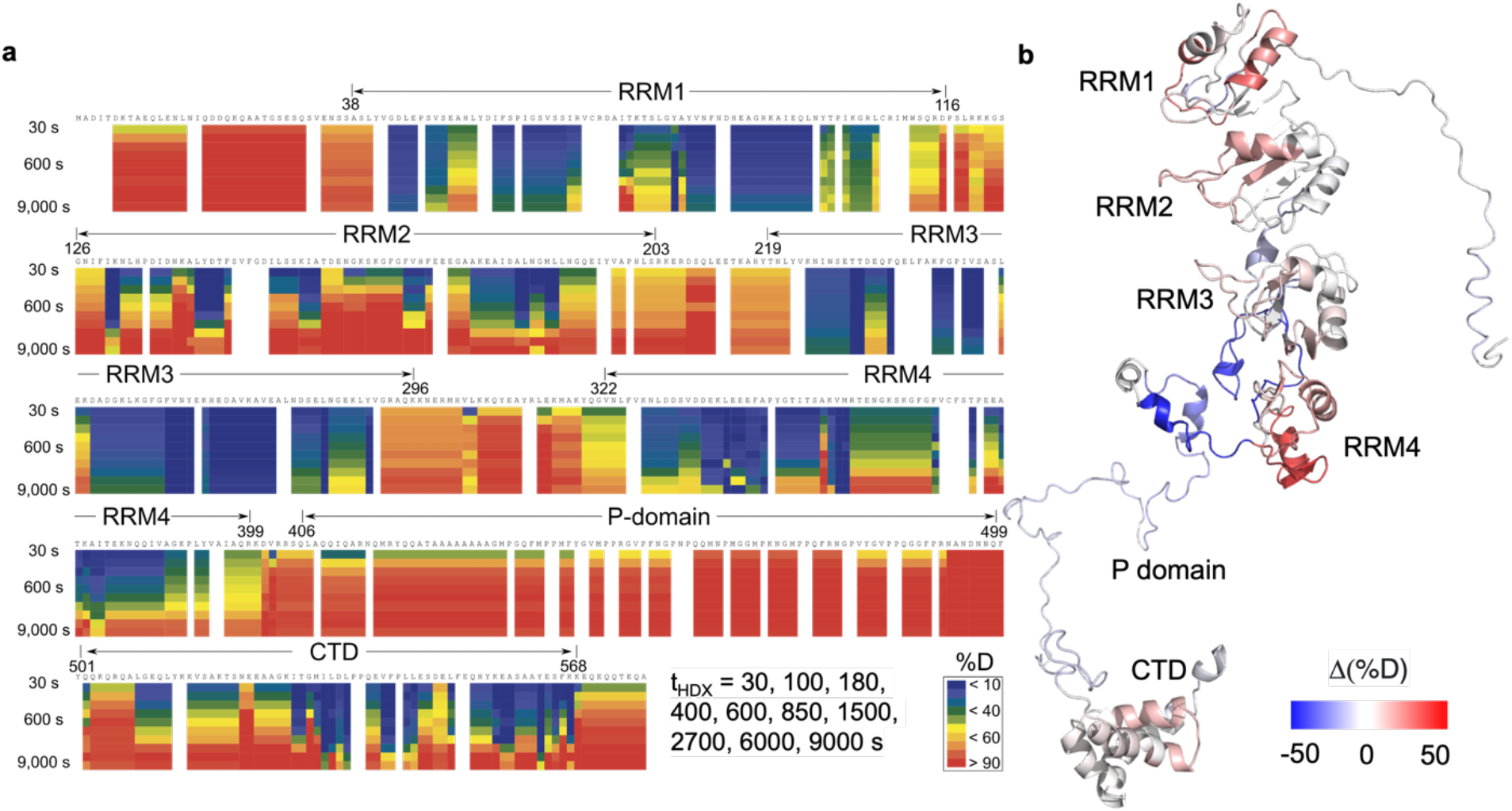
HDX provides structural information of both Pab1 monomer and condensate throughout the sequence. **a**, Heat map of monomeric (soluble) Pab1. Horizontal axis represents Pab1 sequence and vertical axis is increasing HDX timepoints at 30, 100, …, 9,000 s. Color represents the level of deuterium uptake. Boundaries of Pab1’s domains are marked on plot. Plot made with HDExaminer. **b**, *%D* difference between T-condensates and monomer at t_HDX_=100 s (data in Fig. 3a, red) mapped onto Pab1 structure. Structure generated with raptor-X^37^. HDX data were reproducible for 2 bio-replicates (Extended Data Figs 1b-d).

**Fig. 2.**
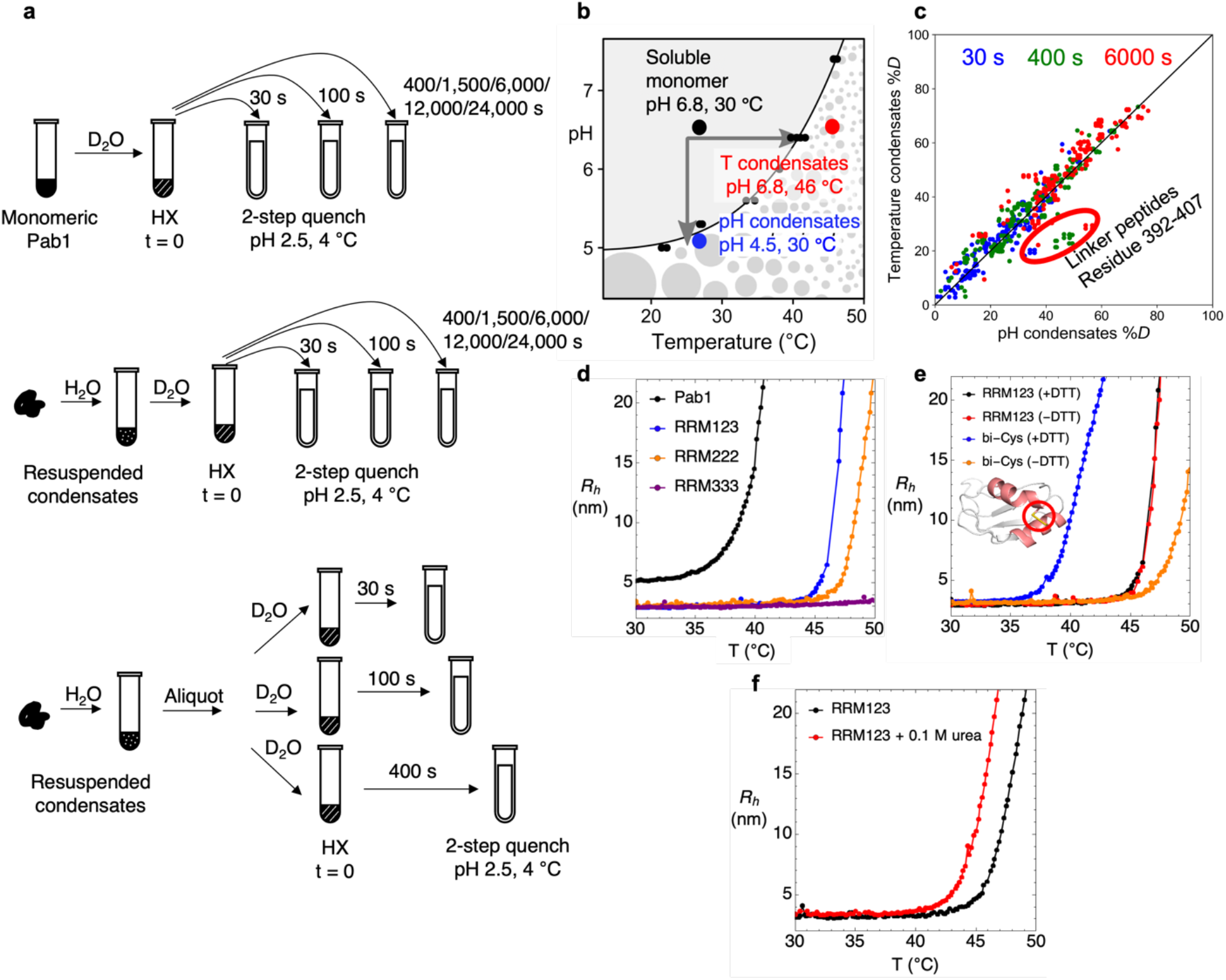
Measurement and HDX labeling of Pab1 condensation induced by thermal and pH changes. **a**, HDX labeling protocols for monomeric Pab1 and condensates. The samples at each time point for each type of condensate were prepared either from a single bulk exchanging sample, or from samples exchanging separately. The two protocols produced consistent results (Extended Data Figs 2c, 2d). **b**, Soluble monomer, temperature-induced condensates and pH-induced condensates were prepared according to Pab1’s phase diagram. Figure is adapted from Riback et al. (2017a)^5^. **c**, Scatter plot comparing the deuterium uptake of pH- and temperature-condensates at t_HDX_=30, 400, 6,000 s. **d**, Condensation of Pab1 (T_cond_ = 39 °C), RRM123 (T_cond_ = 46 °C) and triplet-RRM constructs were measured by DLS. **e**, Condensation of bi-Cys mutant in the absence of 1 mM DTT where a stabilizing disulfide bond is present. The structure of the engineered disulfide bond (I61C, K94C) on RRM1 is shown. **f**, Condensation of RRM123 with or without the presence of 0.1 M urea.

A major challenge for structural studies of biomolecular condensation is that condensation interferes with most solution-based techniques including NMR, which requires rapid molecular tumbling^17^. HDX-MS, however, can probe the hydrogen bond network even in insoluble milieus^18^, all while providing residue-level thermodynamic and structural information^19^. Recent progress in employing crosslinking-MS (XL-MS) to map interactions in the condensates revealed that chaperone HspB8 protects FUS condensates by binding the RRM domain of FUS^20^. In a study of the LCR domain of TDP43, hydroxyl radical footprinting-MS identified methionines involved in forming cross-structures in droplets^21^. While powerful, these two MS-based strategies modify the molecule of interest and provide limited structural or thermodynamic information. In contrast, HDX-MS obtains this information for nearly every residue in a more uniform manner with negligible perturbation^22^. Here we apply HDX-MS to identify the factors that enable Pab1 to transduce physiological cellular stress signals into condensation.

We conducted HDX-MS on three samples, monomeric (soluble) Pab1 (pH 6.8, 30 °C), pH-induced (pH 4.50, 30 °C, 30 min.), and temperature-induced condensates (pH 6.80, 46 °C, 20 min.) (Fig. 2b). Samples were diluted 29-fold into D_2_O buffer (pD_read_ 6.0, on ice) to initiate deuterium labeling. After 30-24,000 seconds of labeling, HDX was quenched at pH 2.5. The condensates then were solubilized with 8 M urea (Fig. 2a) and injected onto LC-MS analysis with an in-line protease column^23,24^. Our peptide map had 99% sequence coverage and an average redundancy of 7 peptides covering each residue, providing a system well-suited for HDX-MS.

For monomeric Pab1, the HDX pattern reflected the boundaries of Pab1’s domains. Peptides from Pab1’s five structured, H-bonded domains exchanged with observed rates, *k*_obs_, considerably slower than the intrinsic (chemical) exchange rate, *k*_chem_, expected for an unstructured protein^19,25^. In contrast, peptides from the disordered regions, including the N- and C-termini, inter-domain linkers, and the IDR P-domain, exchanged with *k*_obs_ at or close to *k*_chem_, consistent with these regions lacking stable H-bonding.

The four RRMs had different exchange behavior indicating different stabilities. For RRM1, 2, and 4, most peptides exchanged 10^3^–10^5^-fold slower than *k*_chem_. The protection factor (PF) ratio, *k*_obs_ /*k*_chem_, corresponds to a stability for each RRM of *RT* ln PF ∼ 4-7 kcal mol^-1^, assuming HDX is occurring in the thermodynamic “EX2” limit (SI)^22^. Many of the peptides from RRM3 exchanged even slower and had less than 50% deuteration even after 24,000 s (Figs. 1a, 3b). Accordingly, we estimate the stability of RRM3 to be greater than 7 kcal mol^-1^, making it the most stable RRM, a point that turns out to influence its relative involvement in condensation.

**Fig. 3.**
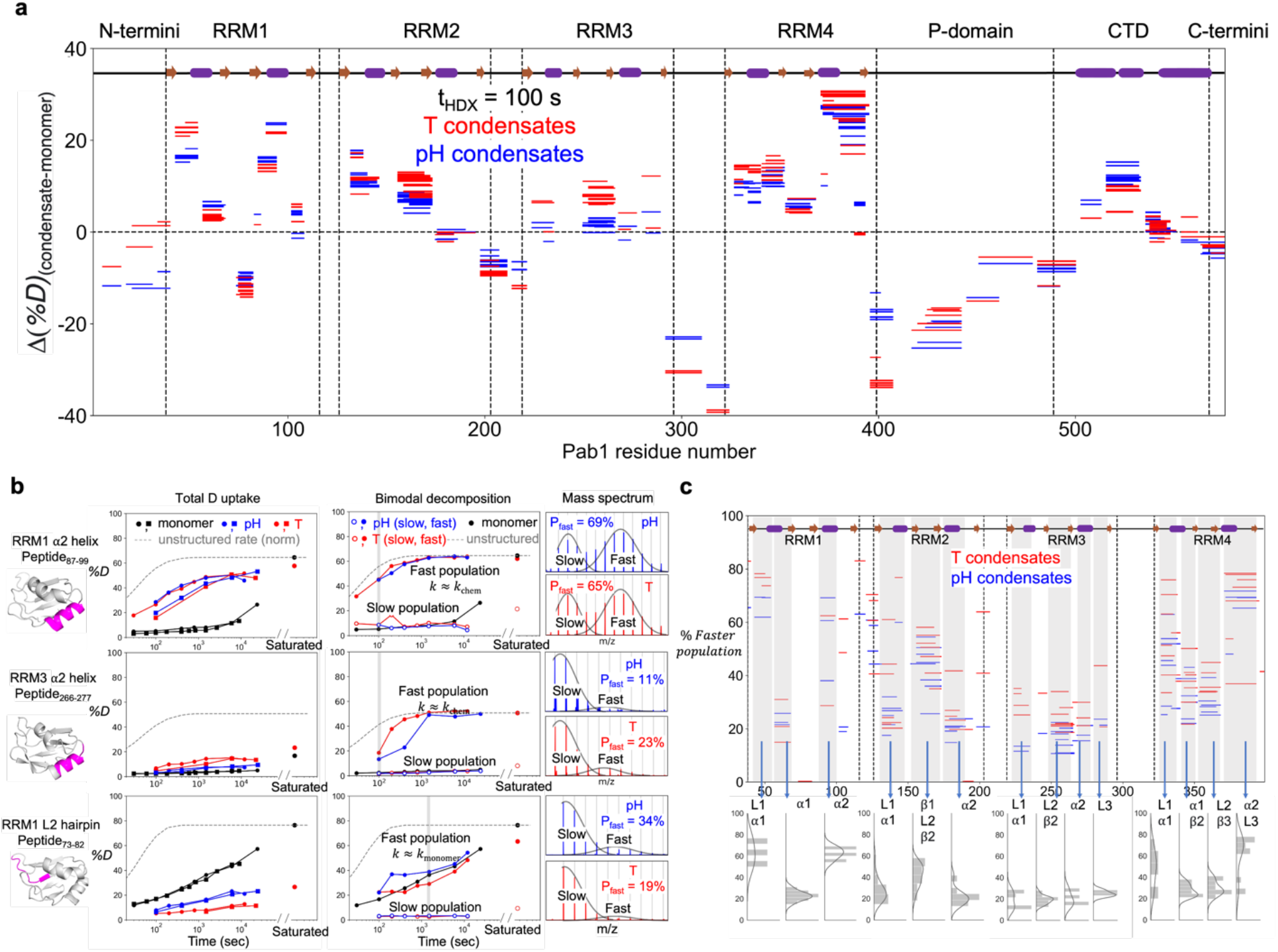
Pab1 condensates exhibited altered HDX behavior and heterogeneity throughout the sequence. **a**, Wood plot comparing HDX of pH- and T-condensates to monomeric Pab1. *%D* differences of both condensates compared to monomeric Pab1 at t_HDX_ = 100 s are plotted against the sequence. **b**, Uptake curves of RRM1 and RRM3 α2-helices, and RRM1 L2 hairpin. Left is the total D uptake curve for the combined fast and slowly exchanging subpopulations where data from two replicates are shown. The middle panel is individual D uptake curves for the faster and the slower subpopulations, obtained using a bimodal fitting procedure. For both plots, the unstructured rate (*k*_chem_) is normalized to the back exchange level. Right is bimodal mass spectrum envelopes indicating two (or more) different subpopulations taken from timepoint 400, 400, 1500 s, respectively, where the two subpopulations are well-separated for optimal fitting. Data from one of the two replicates is shown for simplicity. Corresponding plots for RRM2 and RRM4 α2-helices are presented in Extended Data Fig. 2. **c**, Heterogeneity map of temperature- and pH-induced condensates. Fraction of the faster-exchanging subpopulation for regions of RRMs that become unfolded in the condensates are plotted against the sequence. Peptides from the linker regions are not shown as they are unfolded in the monomeric state. Secondary structures are depicted on top. Density distributions of *%Faster population* for peptides from each secondary structure element are shown at the bottom.

Having established the base-line behavior of the soluble monomer, we then asked how condensation altered the HDX pattern. We observed an extensive change in the HDX pattern for the RRMs with the condensation-altered regions exchanging faster or slower than in the monomer, reflecting both destabilization and stabilization upon condensation, respectively (Figs. 1b, 3a). Less change was observed for the P-domain and C-terminal domain (CTD) (Fig. 3a, Extended Data Fig. 2), consistent with Pab1 condensation being mediated by the RRMs^5^.

Depending on the RRM, destabilized regions included helices, hairpins, and β-strands. Most of the destabilization observed in RRM1 and RRM4 occurred in the α2 helical regions and the L1 hairpin, whereas the most of destabilization observed in RRM2 occurred at the L1 and L2 hairpins instead of the helices (Figs. 1b, 3a, 3b). Some β-strands were also destabilized (Figs. 1b, 3a), which may explain the previous finding that RNA is ejected from Pab1 upon condensation^5^, as the β-strands are responsible for RNA-binding^26^. Compared to the other RRMs, the *%D* uptake of RRM3 was closer to that of the monomeric state (Figs. 1b, 3a), implying condensation had less of an effect on this domain.

Regions stabilized upon condensation included the linkers, P-domain, and some RRM regions (Figs. 1b, 3a). Significantly, the linkers between RRM2-RRM3 and RRM3-RRM4, which were unstructured in monomeric Pab1, became at least 100-fold more protected, suggesting they form stable interactions upon condensation (Fig. 3a). Likewise, HDX was significantly slowed for a region in the P-domain containing eight contiguous alanine residues that may become helical in the condensates (Fig. 3a, Extended Data Fig. 2). Notably, some regions of the RRM even became more protected in the condensates which could be due to a native-like region being stabilized by additional interactions (Figs. 1b, 3a, 3b).

A key feature we observed only in the condensates was widespread structural heterogeneity. The signature of such heterogeneity was numerous peptides having two or more associated peaks in the mass spectrum coming from populations having different deuteration levels (Fig. 3b). In contrast, all peptides in the monomer shifted as a single mass envelope, indicative of a single population. The multimodal HDX data is characteristic of two or more structurally distinct populations that do not interconvert on the labeling timescale, each undergoing EX2 exchange kinetics. However, it could also be due to concerted “EX1” exchange kinetics resulting from a cooperative transition between two states, which is manifested as a peak at lower m/z decreasing in amplitude with a peak at higher m/z correspondingly increasing in amplitude (SI)^22^. In our data, the peak heights remained relatively constant across different time points, implying that the multiple populations were due to EX2 exchange with static heterogeneity, consistent with the non-specific aspect of interactions in biomolecular condensates.

To address the heterogeneity, we fit the bimodal condensate data as a weighted mixture of two populations to extract the time-dependent *%D* buildup curves of each subpopulation and their relative fractions over time (Fig. 3b). Different HDX rates for the subpopulations revealed the unfolding of certain structures and the formation of new contacts. For most fast-exchanging regions, the faster subpopulation exchanged with a rate at or close to *k*_chem_, consistent with the region being unfolded (Fig. 3b, Extended Data Fig. 2). The other major subpopulation exchanged with rates similar to or slower than those observed in the monomeric state (Fig. 3b, Extended Data Fig. 2). The fast-exchanging population fraction ranged from close to zero (RRM3) to greater than 70% (RRM4, α2-helix) (Fig. 3c), reflecting the different unfolding tendencies for different regions of each RRM. For RRM3, the fast-exchanging fraction was under 30% for all RRM3 peptides, consistent with its high stability (Fig. 3c). For a few peptides, both subpopulations exchanged slowly, with one subpopulation exchanging close to the rate observed for soluble Pab1 and the other subpopulation exchanging even slower (Fig. 3b). Again, we attribute this higher protection to a folded region forming additional interactions in the condensate. In summary, Pab1 molecules can adopt multiple, partially unfolded conformations and form a variety of new inter-chain interactions in the condensates.

To investigate how Pab1 integrates signals of thermal and non-thermal (e.g., starvation) stresses into one phase boundary, we compared HDX-MS for both temperature- and pH-induced condensates. We found that their HDX behavior highly similar in both total uptake and heterogeneity (Figs. 2c, 3a, 3b, Extended Data Fig. 1e). For regions with heterogeneity, the averaged *%D* differences between the pH- and T-condensates were 0.3 ± 10 and -0.4 ± 8.3 for the slow and fast exchanging subpopulation, respectively (averaged across all peptides and timepoints, Extended Data Fig. 3). This similarity indicates that Pab1 undergoes similar structural changes therein in response to temperature or starvation-induced acidification (potentially mediated by charge repulsion upon protonation of the histidines, Extended Data Fig. 4), allowing for a unified response with Pab1 as a central sensor^2^.

To investigate whether the disorder observed in the condensates reflects the intrinsic thermal stability of each of the domains, we collected individual ^1^H-^15^N HSQC spectra for RRM1, 2 and 3 at 35 and 45 °C, which are below and above Pab1’s T_cond_ of 39 °C respectively (pH 6.8). At both temperatures, the NMR spectra for the three individual RRM domains remained well-dispersed, the hallmark of a folded protein, and were nearly unchanged with no evidence of any conformation change upon heating (Extended Data Fig. 5a). Furthermore, the peak volumes did not measurably decrease implying all the protein molecules stayed in solution, as determined by a comparison between the area of the methyl and reference compound (Trimethylsilylpropanoic acid, TMSP) peaks (Extended Data Fig. 5b). For RRM1, the most dynamic RRM, the NMR T_1_/T_2_ relaxation measurements, which are sensitive to tumbling rates^17^, were similar at both temperatures (Extended Data Fig. 5c). Further, the relaxation measurements yielded an estimated molecular weight of 9.35 ± 0.42 kDa at 35 °C and 8.54 ± 0.97 kDa at 45 °C, close to RRM1’s molecular weight of 9.6 kDa. Also, in the NMR water saturation-transfer HDX measurements (CLEANEX^27^) at 45 °C, no new peaks appeared that could be associated with rapid HDX and disorder (Extended Data Fig. 5a). Finally, at pH 5.2 (0.2 units below pH_cond_ at 30 °C), RRM1’s spectrum likewise remained similar to its pH 6.8 spectrum (Extended Data Fig. 5d). In summary, these three RRM domains individually remained folded under conditions where Pab1 condenses.

This important finding implies that the disorder observed in the condensates is conditional upon new inter-chain contacts stabilizing the otherwise unstable (higher free energy) partially unfolded forms. In other words, the condensate forms from rare partially unfolded conformations populated enough by stress that they can cluster into a critical nucleus and then grow via diffusion-limited addition of other rare conformations^28,29^. Evidence of the presence of such partially unfolded states that escape detection by standard NMR measurements comes from numerous HDX studies where exchange rates are faster than expected if exchange requires global unfolding^30,31^. Here, HDX occurs from partially unfolded states that have lower free energy than the globally unfolded state. Overall, our findings point to a mechanism that involves otherwise unstable, partially unfolded states forming a network stabilized by inter-RRM interactions within the condensate.

The different levels and patterns of HDX suggested that RRMs participated differently in the condensate. Specifically, RRM3 remained largely folded, whereas RRM1, 2, and 4 exhibited significant local unfolding. We investigated the extent to which each RRM promotes condensation by measuring the T_cond_ of triplet RRM constructs, termed RRM111, RRM222 and RRM333, which retain the native linkers, along with the WT triplet RRM123 (RRM1-RRM2-RRM3) by DLS (Fig. 2d). RRM111 precipitated at room temperature, whereas RRM123 and RRM222 condensed at 46 and 48 °C, respectively. RRM333 remained monomeric with an unchanging R_h_ even up to 50 °C (Fig. 2d). The relative order of T_cond_, RRM111 << RRM123 < RRM222 << RRM333, largely tracks with the extent of HDX seen in each RRMs. These results suggest that each RRM becomes partially unfolded and capable of participating in the condensation process at a different temperature.

A key test of the role of local unfolding in mediating condensation is that suppressing unfolding should suppress condensation. We targeted RRM1’s helices, the most unfolded structures according to HDX-MS. We made a “bi-Cys” variant of RRM123 with a stabilizing disulfide bond inserted between the two helices on RRM1 (I61C, K94C) and lacking the two endogenous cysteines (Fig. 2e), permitting oxidation-dependent stabilization of RRM1. Under reducing conditions where the disulfide bond was absent (1 mM dithiothreitol, DTT), the T_cond_ was considerably lower, 39 versus 48 °C. As a control, the T_cond_ of WT RRM123 was found to have no dependence on DTT. The WT RRM123’s T_cond_ was noticeably higher (46 °C) than the reduced bi-Cys version, presumably because the cysteine substitutions are destabilizing. Regardless, the large increase in T_cond_ upon formation of the disulfide bond indicates that the unfolding of one or both of RRM1’s helices helped trigger condensation in RRM123. In addition, the bi-Cys mutant with the disulfide bond had nearly the same T_cond_ as RRM222 (Extended Data Fig. 1f). This suggests that when RRM1’s partial unfolding is inhibited, the onset of condensation is determined by RRM2, the next domain to undergo partial unfolding.

Inspired by the proposed role of the non-specific weak interactions in phase separation^8–11^, we evaluated whether each molecule must contain at least one partially unfolded RRM to participate or whether weak interactions are sufficient for an RRM to participate in condensation. We examined whether triplet RRM constructs (RRM123, RRM222 and RRM333, T_cond_ = 46, 48 and >50 °C, respectively) would co-condense with Pab1 at a temperature (42 °C or 46 °C) above Pab1’s T_cond_ (39 °C) but below the triplet’s T_cond_. The amount of material that co-condensed was identified using SDS-PAGE from the fraction in the pellet after centrifugation at 158,000 g for 10 min. When the triplets were mixed with Pab1 at 42 °C, there was little (RRM123, 222) to no measurable (RRM333) amount of triplet in the condensate (Extended Data Figs. 6a-c). At 46 °C (at or below T_cond_ of the triplets), the majority of RRM123 (>50%) and almost all the RRM333 molecules (>90%) remained in the supernatant (Extended Data Figs. 6a, 6c). The majority (>50%) of the RRM222 molecules, however, co-condensed (Extended Data Fig. 6b). We attribute this difference to RRM222 having three domains on the verge of co-condensing, which helped increase the number of interactions (i.e., valency) that can promote its participation in the condensate. In addition, the interactions between RRMs can be heterotypic according to the co-condensing measurement between RRM113 and RRM222 (Extended Data Fig. 6d). In summary, given the difference between the extent of co-condensing at 42 and 46 °C, and the lack of co-condensing for RRM333, we conclude that domain partial unfolding is the major requirement for participation in the condensation process.

Further supporting our conclusion that partial unfolding is a necessary step, we found that condensation was promoted by low concentrations of denaturant. The addition of 0.1 M urea reduced T_cond_ by 2 °C (Fig. 2f), whereas 1 M added urea abolished condensation even though circular dichroism (CD) spectroscopy indicates no loss of secondary structure (Extended Data Fig. 7). The contrasting effects of denaturant support a model where Pab1’s condensation involves an initial partial unfolding step followed by the formation of a network of interchain interactions. The unfolding step is promoted by denaturant, but the subsequent condensate-promoting interactions are hindered, with the net effect favoring condensation only at low concentrations of denaturant.

Small-angle X-ray scattering (SAXS) measurements found that monomers can be modeled as an ensemble of self-avoiding, folded but non-interacting domains connected by flexible linkers (Extended Data Fig. 8). The experimental pair-distance distribution function, P(r), was similar to that for a non-interacting ensemble, especially as compared to an extended or compact conformation. In addition, the P(r) was largely unchanged when the inter-domain interactions were reduced, e.g., going from 50 to 350 mM NaCl or adding 0.5 M urea. These results support our modeling of the ensemble as non-interacting domains and argue against an auto-inhibition model where folded RRMs on the same molecule interact to generate a compact state that inhibits the inter-molecular interactions necessary for condensation.

Based on our findings, we propose a *sequential activation* model for Pab1’s condensation (Fig. 4a). As temperature increases, each of the RRMs becomes sequentially activated, in this case reflecting different thresholds for local unfolding. Our NMR studies found that isolated RRM domains remained folded at temperatures above Pab1’s T_cond_. Therefore, activated RRMs need additional interactions with other regions to remain partially unfolded. The heterogeneity in the HDX data suggests a high degree of structural diversity with a variable set of contacts in the condensates. Additional interactions are facilitated by an increase in local concentration, which results in the chain optimizing the intermolecular interactions in a maturation or hardening process. This process stabilizes the condensate^9,32^ to the point where it cannot be reversed by cooling (Extended Data Fig. 1a).

**Fig. 4.**
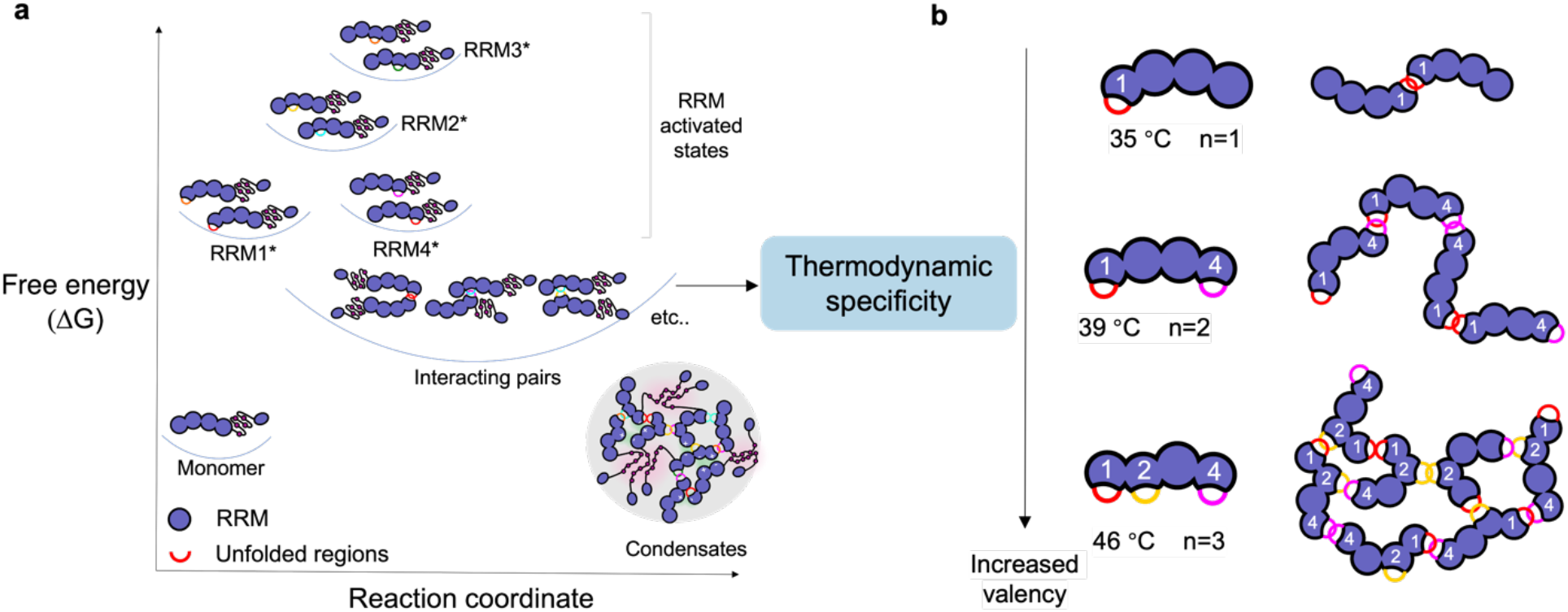
Sequential activation controls Pab1’s condensation, and RRMs have thermodynamic specificity. **a**, When the temperature is increased, each RRM becomes sequentially activated, reflecting its proclivity to partially unfold and associate with other molecules. Activated RRMs have free energies higher than that of the monomeric state. Different colors of unfolded regions represent the heterogeneity that each molecule can have different regions unfolded. An activated RRM can interact with likewise activated RRMs on other chains and facilitate the formation of a highly interconnected, stable network. **b**, Thermodynamic specificity suggests a change in condensate morphology with increasing temperature as the interaction valency (i.e. number of activated RRMs) increases.

Because the RRMs have different activation temperatures and the ability to form stable contacts requires domain activation, the RRMs will tend to form contacts primarily with other RRMs having similar activation temperatures. We term this partner selection *thermodynamic specificity*. That is, condensation does not depend upon the presence of the highly complementary interface that one expects for biomolecular binding. Rather, multivalent interactions between activated RRM domains determine the contact network. On a chain with multiple classes of RRM, the activation of additional domains at higher temperatures increases the valency. This increased valency lowers the percolation threshold and increases the likelihood of forming a system-spanning network^33,34^. In principle, the valency increase with temperature should lead to an increase in condensate stability and change in morphology, successively changing the ensemble from dimers to 1D chains and then to a 3D network (Fig. 4b).

Our study finds a unifying molecular portrait of Pab1 condensation triggered by different stresses with stress-induced local unfolding and domain activation playing a major role. Even though mediated through the folded RRM domains rather than the IDR P-domain^5^, Pab1 condensation conforms to the standard view that disorder and multivalent interactions (usually mediated by LCR/IDR for other condensing proteins) underlie Pab1 condensation. It remains to be determined whether RRMs are especially capable of forming condensates.

Our findings also reconcile different views in the historical understanding of thermal stress. A long-standing model proposes thermally induced proteotoxicity resulting from misfolding and aggregation combated by molecular chaperones^35,36^, yet recent results demonstrate that thermally induced condensates of Pab1 are adaptive and dispersed by chaperones with extraordinary efficiency^5,7,14^. Our current findings provide strong support for a merged paradigm in which Pab1, and presumably other stress-condensing molecules, undergo limited unfolding in order to condense, simplifying the molecular dispersal process mediated by chaperones and promoting rapid adaptation as a consequence.

## Methods

### Condensates preparation

Condensates were prepared by subjecting a 60 μM Pab1 stock to the condensing condition (pH 6.80, 46 °C for 20 minutes for T-condensates; pH 4.50, 30 °C for 30 minutes for pH-condensates). After treatment, the samples were centrifuged at 15,800 g for 10 min. The supernatant was removed, and the pelleted condensates were washed 2x by buffer under the same centrifugation condition.

### HDX labeling

We prepared two HDX replicates using proteins from cells grown separately (2 bio-replicates). HDX was initiated by diluting samples 29-fold into D_2_O buffer (50 mM sodium phosphate, 100 mM NaCl, pD_read_ 6.00) on ice. For each state and each replicate, samples at HDX timepoints of 30 s, 400 s, 1,500 s, 6,000 s, 12,000 s, and 24,000 s were taken. Additional points at t_HDX_ = 10 s, 200 s, 800 s were taken for some states/replicates.

At each timepoint, a quench buffer containing 600 mM Glycine, pH 2.5 and 8 M urea was added at 1:1 ratio to stop HDX and dissolve the condensates. The quenched sample was further diluted with buffer containing 600 mM Glycine, pH 2.5 at 2:1 ratio to arrive at a final urea concentration of 2.67 M to avoid denaturing the protease. (Fig. 2a).

For condensate replicate 1, condensates were resuspended and aliquoted. Then D_2_O buffer was added to each aliquot separately. At corresponding HDX timepoints, quench buffer containing 600 mM Glycine, pH 2.5 and 8 M urea was added at 1:1 ratio to quench HDX and dissolve the condensates. The quenched sample was further diluted with buffer containing 600 mM Glycine, pH 2.5 at 2:1 ratio to arrive at a final urea concentration of 2.67 M.

For replicate 2, condensates were resuspended with a minimum amount of H_2_O buffer and then diluted into D_2_O buffer as a bulk HDX reaction, then individual aliquots were removed and quenched at each timepoints using the same protocol as for the monomeric Pab1.

All-D (fully deuterated) control samples were labeled in 50 mM sodium phosphate, 100 mM NaCl, pD 8, using the same protocol for other data points. For some stable regions including the α2-helix of RRM3 and the slow populations for most peptides in the condensates, the control samples were not fully deuterated. To avoid confusion, we term these controls as “saturated” instead of “all-D”.

### LC-MS

LC-MS measurements were done as described in Zmyslowski et al., (2022)^38^.

### HDX-MS data analysis

For assignment, MS/MS data was searched against a sequence database containing sequences of Pab1, protease, and other proteins running on the LC-MS system using SearchGUI software (CompOmics Group). Search settings: unspecific cleavage, precursor charge 1-8, isotopes 0-1, precursor m/z tolerance 10.0 ppm, fragment m/z tolerance 0.5 Da, no post-translational modifications, peptide length 6-30. The result of the search was imported into PeptideShaker (CompOmics Group) and further processed by EXMS2 (http://hx2.med.upenn.edu/EXms/) to generate a peptide list.

For HDX analysis, MS1 data together with peptide list were imported into HDExaminer 3.1 (Sierra Analytics) to fit peptide isotope distributions. For condensates where some peptides have distinct bimodal mass envelopes, the bimodal fitting option in the software was used to determine the mass centroids of two individual populations. Downstream analysis and plotting were performed with a Jupyter Notebook. HDX data exported from HDExaminer was filtered by confidence score >= 0.88 for analysis and plotting. For bimodality/heterogeneity analysis, a filter of peptide length >=9 was applied to ensure enough separation of the spectra of two populations for a good bimodal fitting quality.

### DLS

DLS measurements were done as described in Riback et al., (2017a)^5^. Measurements were performed in DynaPro NanoStar (Wyatt Technology). Measurements were performed as a slow temperature ramp at 0.25 °C min^-1^ continuously from 25 °C or 30 °C. All experiments unless noted, were performed at 15 μM concentration in 20 mM HEPES, pH 6.4 with 150 mM KCl, 2.5 mM MgCl_2_, and either in the absence or presence of 1 mM DTT. T_cond_ is characterized as the temperature where R_h_ is twice the pre-transition value (R_h_ ∼ 5 nm for Pab1, ∼ 3 nm for RRM123).

### Protein expression and purification

Protein expression and purification protocols were adapted with modification from Riback et al., (2017a)^5^. BL21 *E. coli* cells transformed with an expression plasmid for N-terminally 8xHis-tagged protein constructs were grown in LB at 37 °C until the optical density at 600 nm (OD600) reached between 0.6 and 0.7. The flask was cooled down at room temperature for 30 minutes before being transferred to a 30 °C incubator. IPTG was added to a final concentration of 0.2 mM to induce protein expression. Bi-Cys mutant protein was expressed in Shuffle T7 competent *E. coli* cells (NEB, cat# C3026J) and cells were grown at 30 °C instead of 37 °C.

Cells were harvested after 4 hours of induction and lysed by sonication on ice in buffer containing 20 mM HEPES, pH 6.7, 150 mM KCl, 5 mM imidazole, 1 mM PMSF and 0.1 % Triton. Lysate was clarified at 13,000 g for 30 minutes and loaded onto a 5 mL HiTrap chelating HP column (Cytiva 17-0409) on a Bio-Rad FPLC system. Bound protein was eluted with an imidazole gradient. Fractions containing the target protein were pooled with -mercaptoethanol and TEV protease and dialyzed into 20 mM HEPES pH 6.7, 150 mM KCl, 10% glycerol, to remove N-terminal TEV tags. TEV-cut fractions were pooled and loaded into two 1 mL HiTrap heparin HP column (Cytiva 17040601) for removal of nucleic acid contaminants with elution over a KCl gradient. Protein concentration was measured by absorbance at 280 nm.

### NMR

NMR data were acquired either on Bruker AVANCE IIIHD 600 or a Bruker AVANCE III 500 NMR spectrometer equipped with a room temperature TXI probe with Topspin 3. Samples were exchanged to buffer at indicated pH (20 mM HEPES at pH 6.8, 150 mM KCl, 2.5 mM MgCl_2_, 2 mM TCEP or 50 mM sodium acetate at pH 5.2, 100 mM KCl, 2 mM TCEP) and concentrated to > 150 μM. The mixing time for CLEANEX-PM measurements was 200 milliseconds. Data were inspected in Topspin and analyzed and plotted using NMRviewJ.

### Total-Soluble-Pellet (TSP) assay and co-condensing measurements

TSP assay for determining the protein fraction in the condensate was adapted from Riback et al., (2017a)^5^. Concentrated protein stocks were diluted into a buffer containing 20 mM HEPES, pH 6.4 with 150 mM KCl, 2.5 mM MgCl_2_ to a final concentration of 10 μM. For co-condensing experiments, both proteins were at a concentration of 10 μM. The sample was incubated at 42 °C for 20 min or 46 °C for 10 min followed by centrifugation at 15,800 g for 10 minutes at 10 °C. Supernatant was collected as the soluble fraction sample. Buffer was added to the pellet to wash out residual supernatant and the sample was centrifuged again under the same condition. After removing the supernatant, the pellet was resuspended in buffer as the pellet fraction sample. Total (T), soluble (S), and pellet (P) fractions are analyzed by SDS-PAGE.

### Small-angle X-ray scattering

SAXS data were collected at the BioCAT beamline at the Advanced Photon Source (Argonne National Lab) using an in-line SEC-SAXS protocol adapted from Riback et al., (2017b)^39^. A concentrated protein sample (∼3 mg/mL) was injected onto an SEC system with a GE Lifesciences Superdex-200 size exclusion column running in 20 mM HEPES, pH 7.4, 150 mM KCl, 2 mM DTT at room temperature. Data were collected and analyzed using RAW^40^ and GNOM^41^ with the R_g_ values obtained from the pair-distance distribution function, P(r), that were calculated using an indirect Fourier transform of the scattering data. The simulated conformational ensembles were created using the *Upside* molecular dynamics algorithm^42–44^.

## Acknowledgments

We thank I. Gagnon, A. Zmyslowski, and M. Baxa for assistance in MS and protein expression, and H. Yoo, H. Glauninger, and J. Bard for useful discussions. This work is supported by NIH grants GM126547 (DAD, TRS), GM055694 (TRS), GM127406 (DAD), GM144278 (DAD), T32GM007183 (JSh). Use of the Advanced Photon Source, an Office of Science User Facility, operated for the Department of Energy (DOE) Office of Science by Argonne National Laboratory, was supported by the DOE under Contract DEAC02-06CH11357 and a grant from the US Army Research Office (W911NF-14-1-0411, DAD).

## Author contributions

R.C., J.A.R., D.A.D. and T.R.S. designed the research; R.C., D.K., J.S., J.A.R. and T.R.S. performed the experiments and analyzed data; R.C., T.R.S., D.A.D., J.A.R. and D.K. wrote the manuscript.

## Additional information

Supplementary information is available for this paper. Correspondence and requests for materials should be addressed to.

## Extended data figure legends

**Extended Data Fig. 1.**
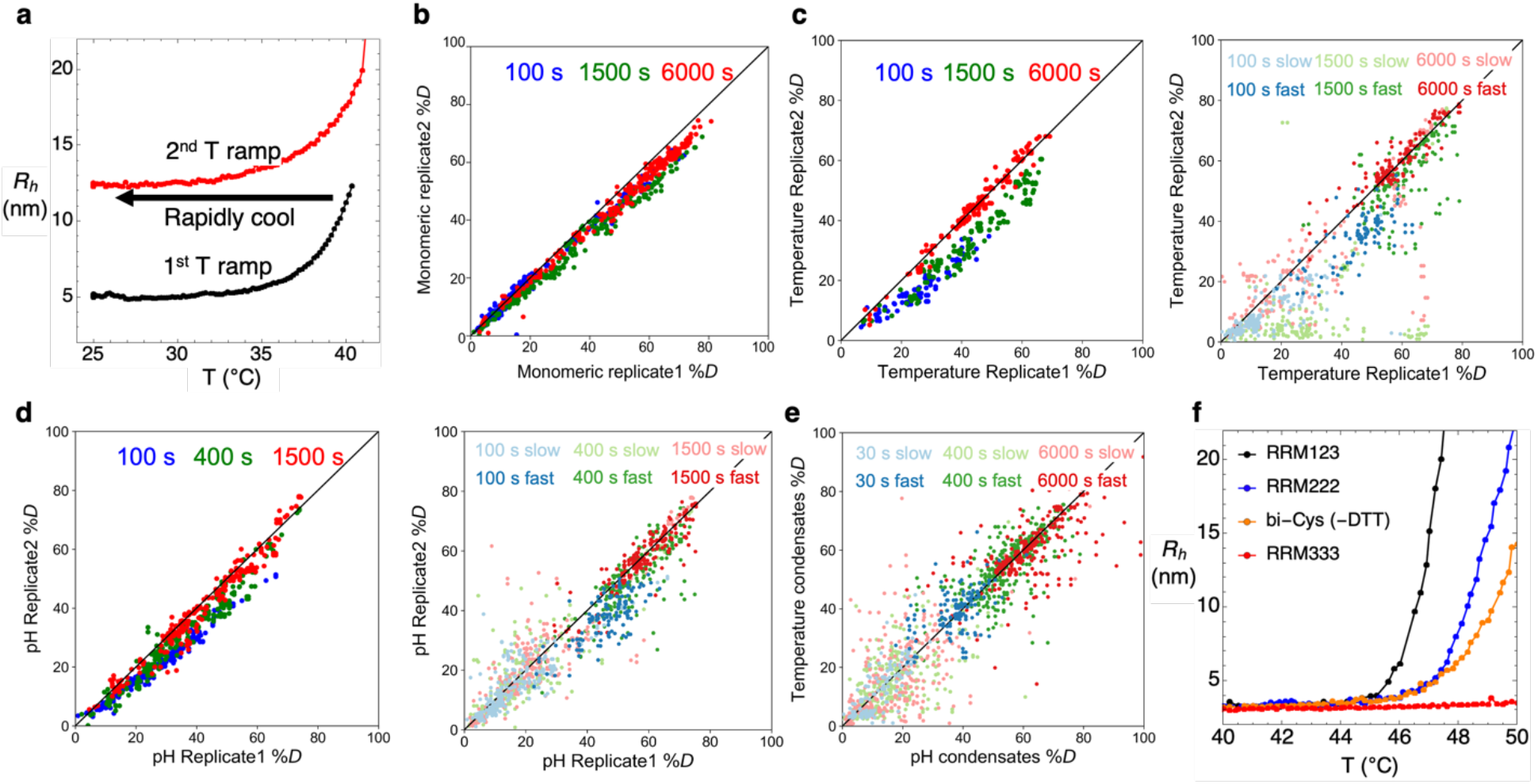
Irreversibility of condensation and reproducibility of HDX. **a**, Pab1 condensation is irreversible by cooling and pre-formed condensates do not nucleate condensation at temperatures below T_cond_. Sample was heated to the onset of condensation (at 40 °C, 1 °C above T_cond_ = 39 °C) and then cooled back to 25 °C for 20 min. The R_h_ level, 11.5 nm, did not change during cooling. The sample was then reheated and the same temperature-dependent profile in R_h_ as observed in the initial heat ramp. **b**, Comparing *D%* of two soluble/monomeric replicates at 3 different timepoints. **c**, Comparing *D%* of two temperature-condensate replicates. Left is *D%* difference of overall uptake (sum of the fast and slower exchanging subpopulations for regions with heterogeneity); right is *D%* difference of faster or slower exchanging subpopulation. The reduced correlation is likely a result of the added complication of the bimodal decomposition into fast and slowly exchanging subpopulations. **d**, Comparing *D%* of two pH-condensate replicates. Similar to figure c, left is *D%* difference of overall uptake and right is *D%* difference of the faster or slower exchanging subpopulation. **e**, *D%* difference of faster or slower exchanging subpopulation for pH- and temperature-condensates. The corresponding plot showing the overall uptake is presented in Fig. 2c. **f**, Bi-Cys mutant with the disulfide bond has nearly the same T_cond_ as RRM222.

**Extended Data Fig. 2.**
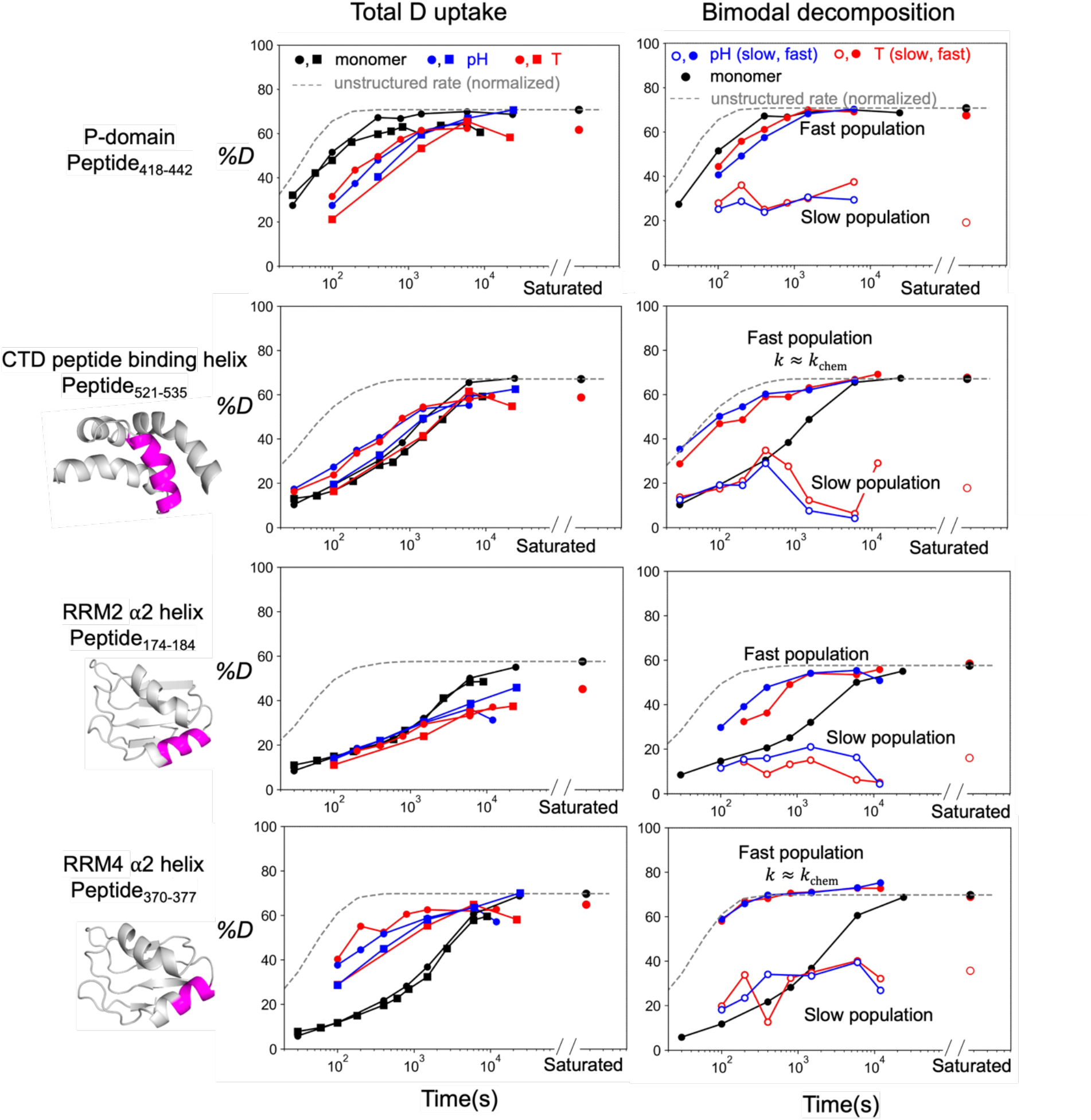
Uptake curves for peptides from the P-domain, CTD, and α2 helices in RRM2 and RRM4. Left is total *%D* uptake curve for the combined fast and slowly exchanging subpopulations from 2 replicates were shown. Right is *%D* uptake curves for the faster and the slower subpopulations obtained using a bimodal fitting procedure. HDX of Peptide_418-442_ from P-domain was slowed for half of the molecules. CTD’s helix 2 (Peptide_521-535_, part of the CTD’s peptide binding site^45^) was unfolded for the majority of the molecules, which potentially led to a disruption of the hydrophobic core and destabilization of other CTD regions (Fig. 3a).

**Extended Data Fig. 3.**
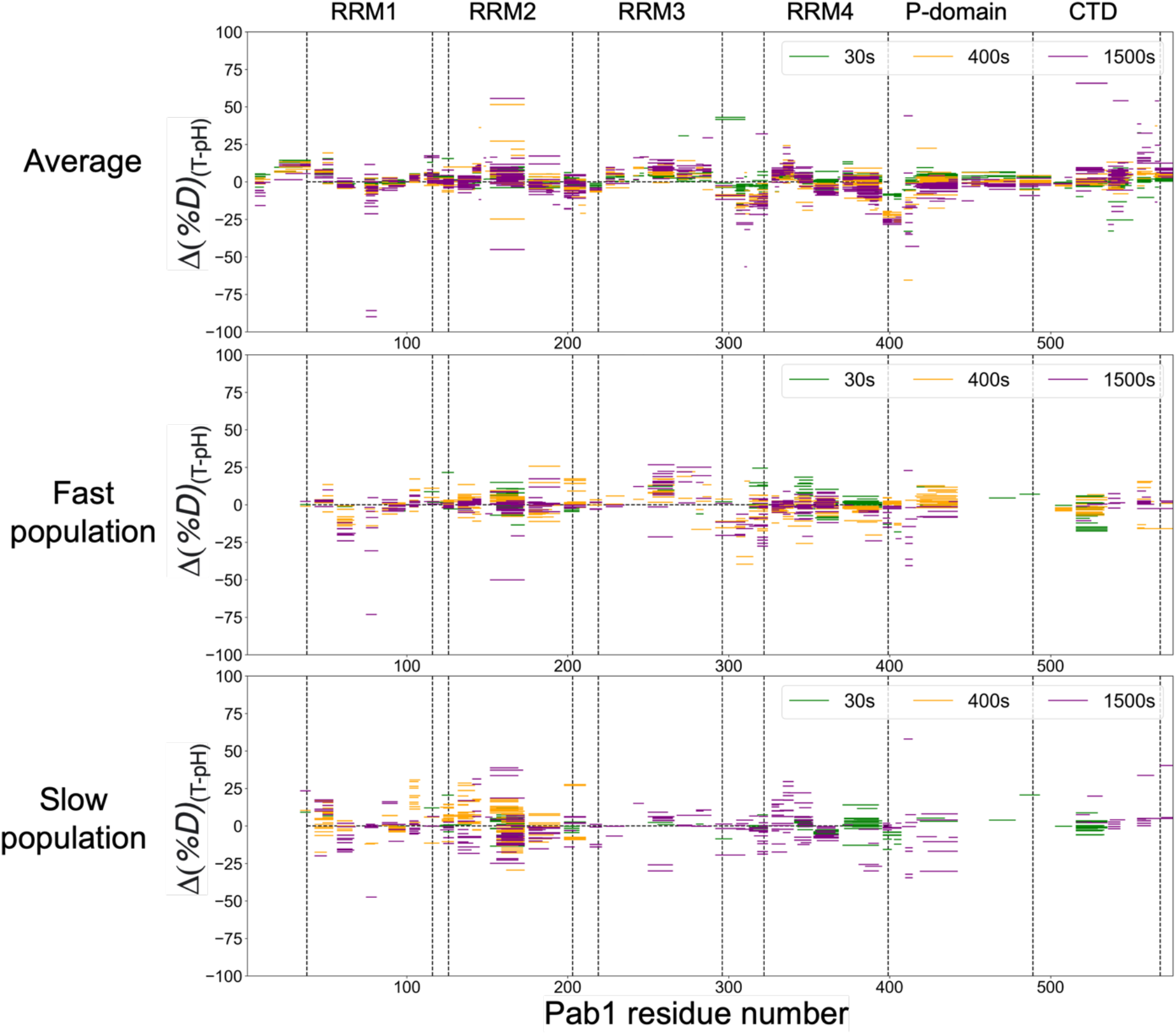
*%D* difference between temperature- and pH-induced condensates at 3 exchange times. **a**, *%D* difference of total uptake (combining the fast and slow exchanging subpopulations for regions with heterogeneity). **b**, *%D* difference of the faster exchanging subpopulation. **c**, *%D* difference of the slower exchanging subpopulation.

**Extended Data Fig. 4.**
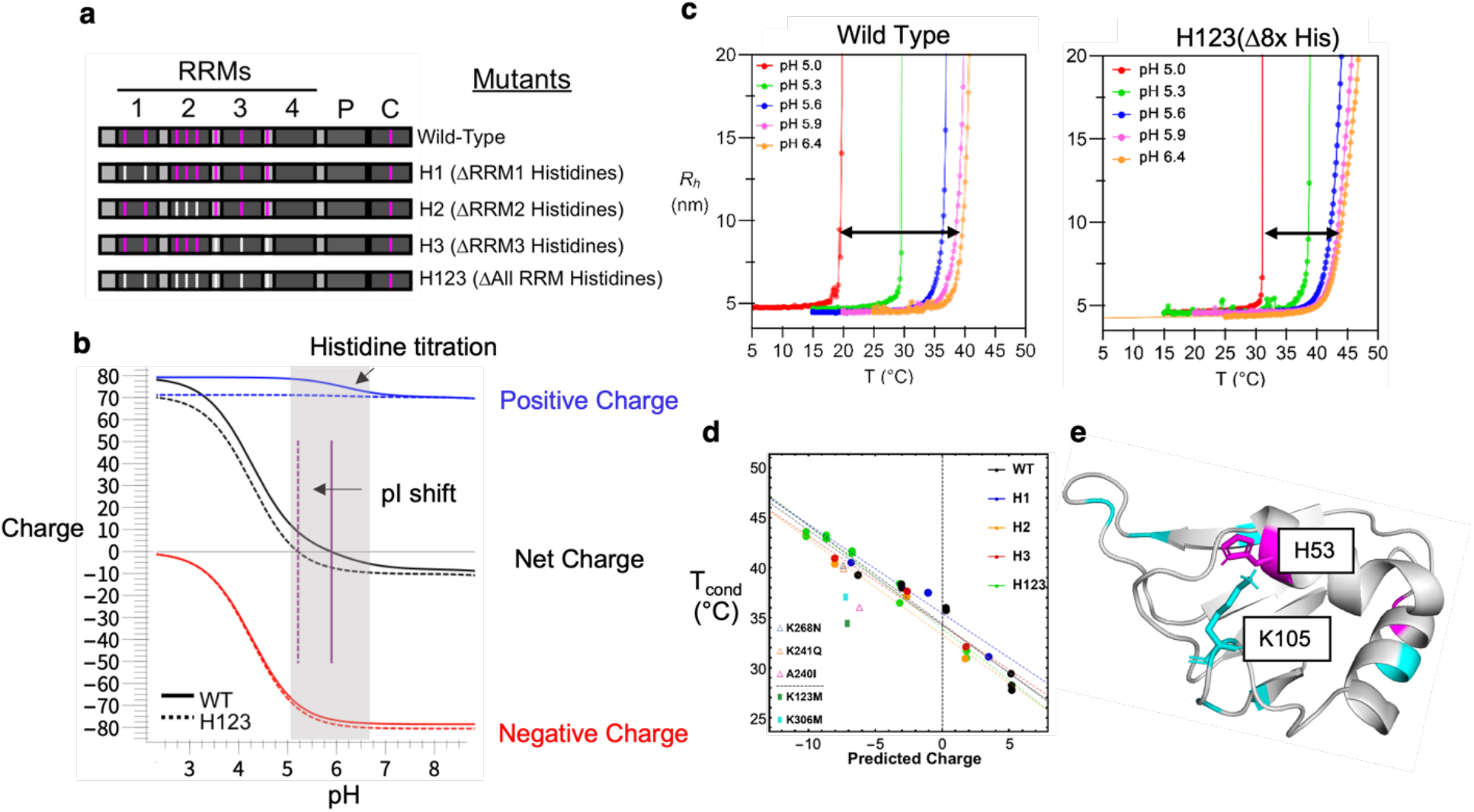
pH sensitivity of Pab1 condensation and the histidines. **a**, Histidines on RRMs and the histidine knockout mutants where histidines from each RRM (H1, H2, H3) or all RRMs (H123) are substituted with the next most common amino acid in the Pab1 sequence alignment^5^. **b**, pH titration curve of WT Pab1 and H123 mutant. Histidine titration overlaps with Pab1’s condensing pH. **c**, DLS of H123 mutant and WT Pab1 at different pHs. Condensation of the H123 construct had a reduced pH sensitivity, with T_cond_ decreasing by 12 °C when pH was reduced from 6.4 to 5.0, whereas WT Pab1 experienced a 20 °C decrease. This decrease in sensitivity implies that histidines are partly responsible for Pab1’s response to acidification. **d**, Predicted charges of Pab1’s mutants correlate with their T_cond_ at different pHs. Non-histidine mutants (K241Q, K268N, K123M, K306M, A240I) deviate from the line, indicating that Pab1 has different sensitivities to different charges. A previous phase separation study investigated a broader range of net charge than ours and observed that the saturation concentration (c_sat_) was the lowest around charge neutrality^46^. **e**, Histidines (magenta) and other positively charge residues (lysine and arginine, colored in cyan) on RRM1. A few histidines on the RRMs are adjacent (< 8 Å) to arginine or lysine residues. Specifically, the positive charge on H53 and K105 are estimated to be 3.9 Å apart, providing a electrostatic repulsion when the histidine is protonated. Upon acidification, histidines become protonated and the destabilizing Coulombic repulsion may cause unfolding of the region and subsequent condensation. Nevertheless, the titration of histidine does not fully explain Pab1’s pH response. Although RRM4 has no histidines, pH-induced condensates still show a similar HDX pattern and comparable level of unfolding as observed with the temperature-induced condensates.

**Extended Data Fig. 5.**
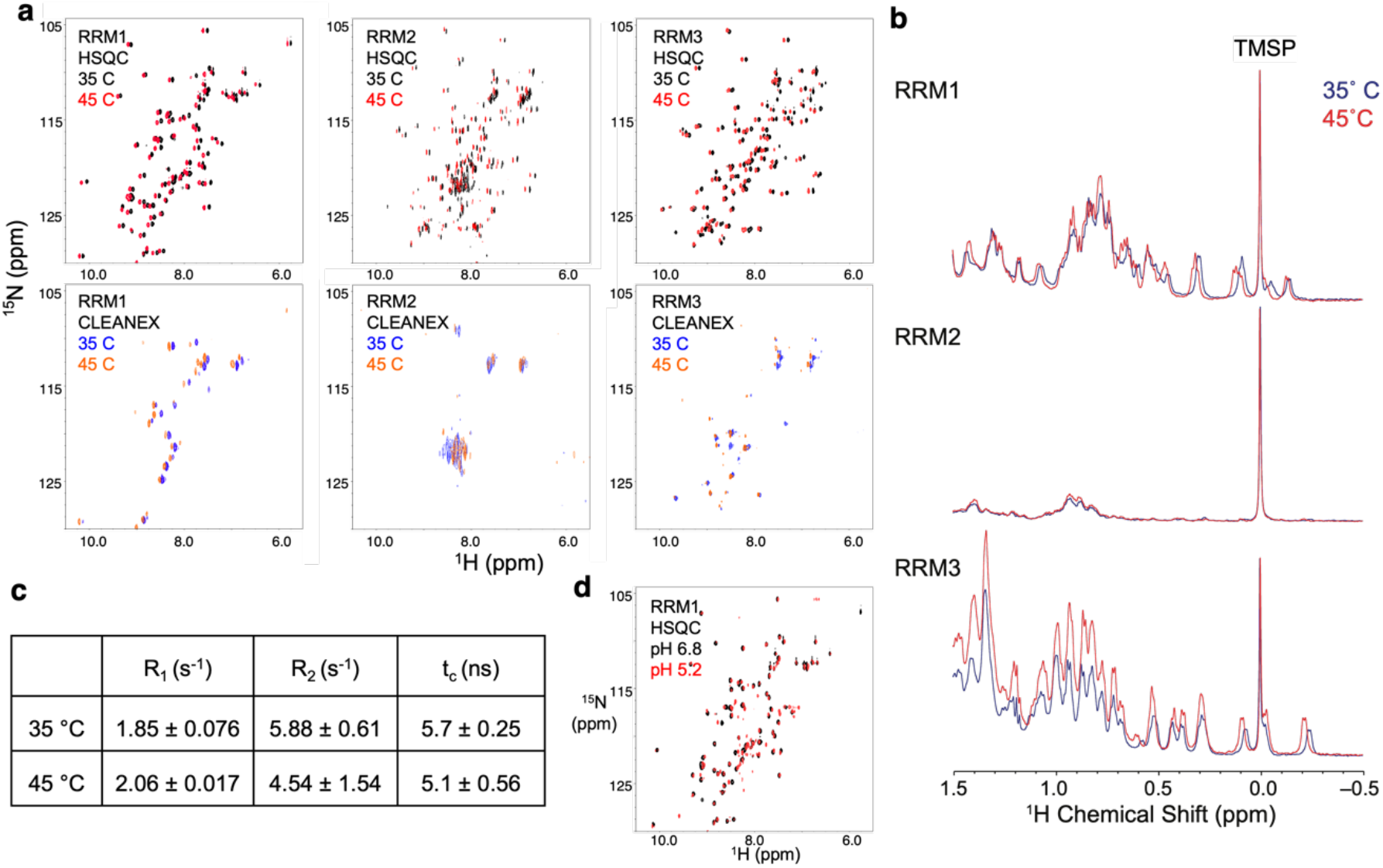
NMR indicates that individual RRMs remain well-folded above Pab1 T_cond_ or below Pab1 pH_cond_. **a**, ^1^H^15^N NMR HSQC spectra (upper) of RRM1, 2 and 3, at 35 and 45 °C, pH 6.8, and corresponding water saturation transfer (CLEANEX^27^) spectra (lower) at the same conditions. **b**, Overlaid ^1^H spectrum at 35 and 45 °C comparing methyl peaks and the reference compound. **c**, Relaxation measurements of RRM1 at 35 and 45 °C, pH 6.8. **d**, Overlaid _1_H^15^N NMR HSQC spectrum of RRM1 at pH 6.8 and pH 5.2, 30 °C.

**Extended Data Fig. 6.**
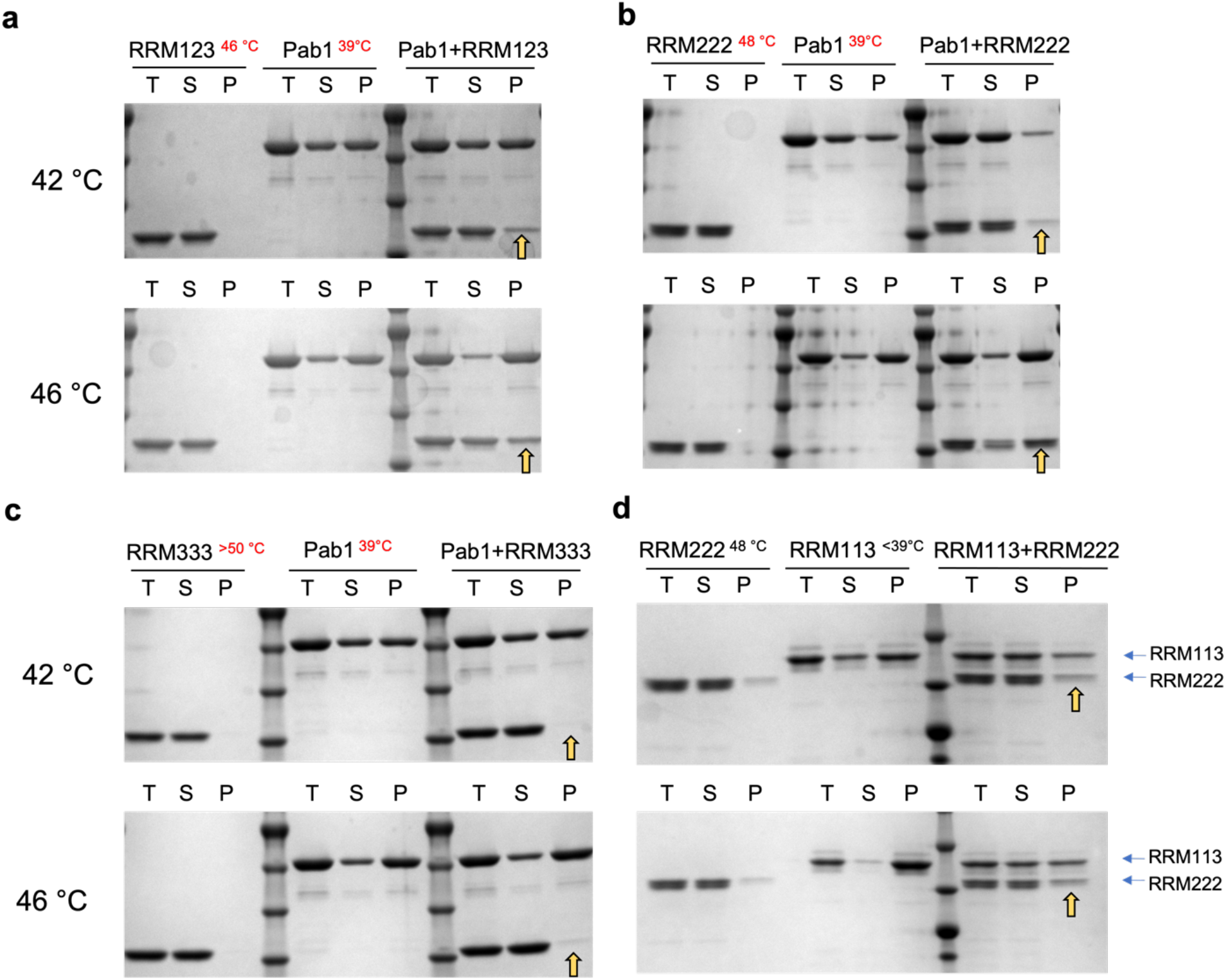
Co-condensing experiments at 42 °C and 46 °C. **a-c**, Co-condensing with Pab1 for RRM123, RRM222 or RRM333. **d**, RRM222 co-condensing with RRM113 at 42 °C (upper panel) or 46 °C. Co-condensing band position is indicated with a yellow arrow. T_cond_ of constructs are provided (superscript). At both 42 or 46°C, a minority of molecules of RRM222 co-condensed into the pellet lane, and RRM113’s condensation was reduced. This result indicates that heterotypic interactions can occur between RRM113 and RRM222, but in a manner that inhibits RRM113’s condensation.

**Extended Data Fig. 7.**
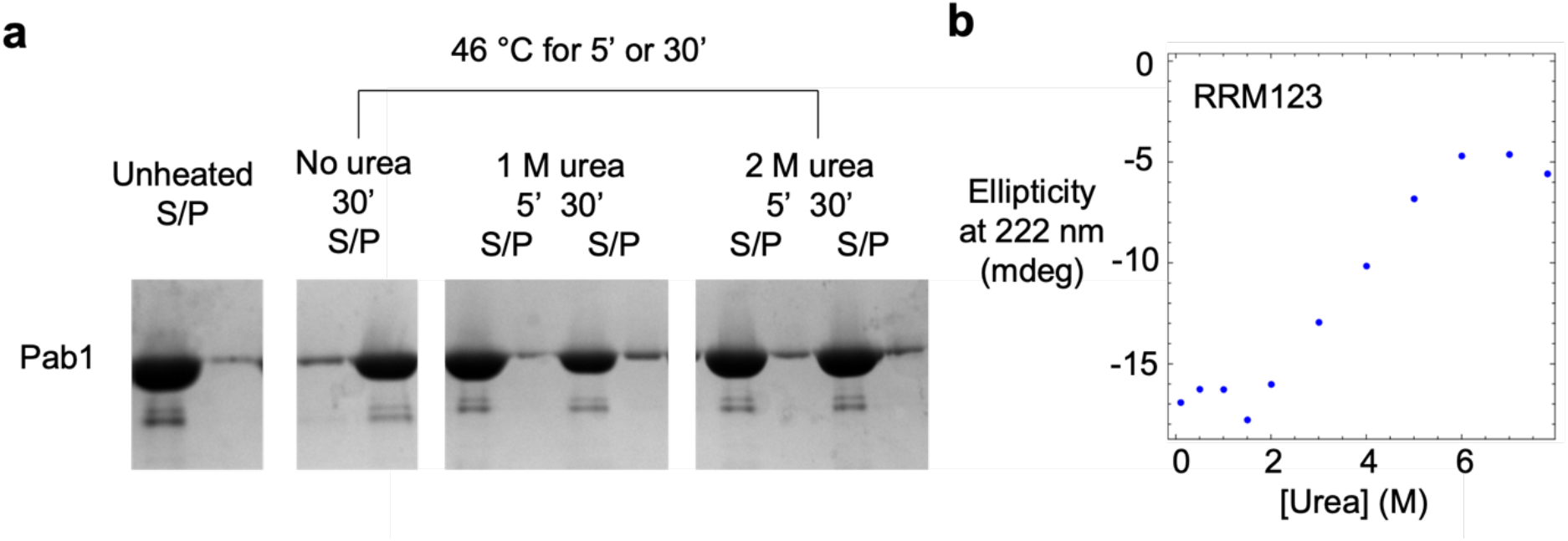
1 M urea abolishes Pab1 condensation. **a**, Pab1’s condensation is inhibited by 1 M urea, analyzed by TSP. The gel lanes are from the same experiment where samples were loaded onto two gels which were treated in parallel. The uncropped gel picture is available in supplementary figures. **b**, Circular dichroism of RRM123 indicates no loss of secondary structure with 1 M urea. Ellipticity at 222 nm is plotted.

**Extended Data Fig. 8.**
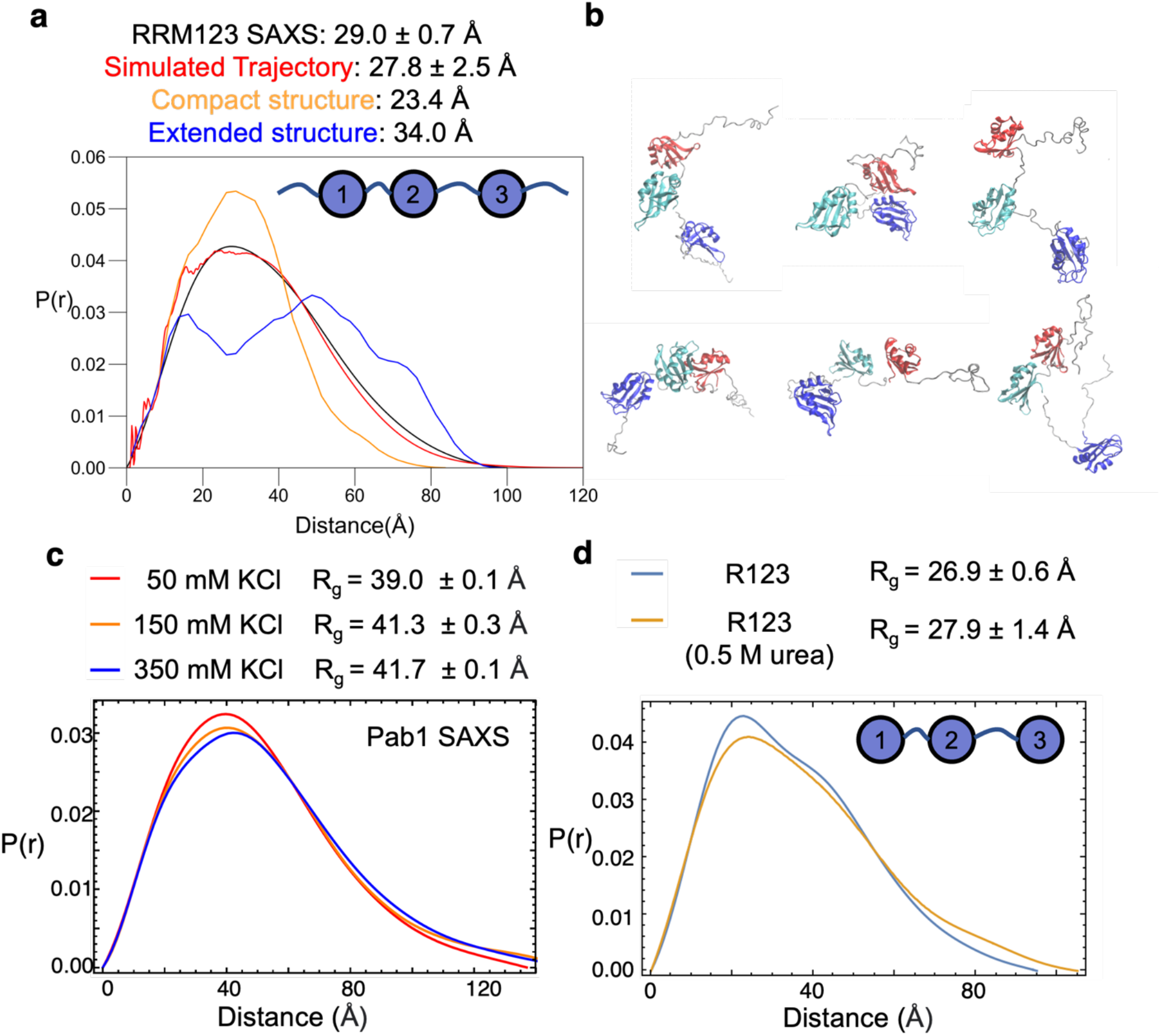
SAXS studies find that the RRMs are not in a compact, auto-inhibited conformational ensemble prior to condensation. **a**, The experimental P(r) for the RRM123 construct having the three amino-terminal RRM domains, is very similar to a P(r) for a simulated conformational ensemble having folded but self-avoiding, non-interacting RRM domains connected by flexible linkers. The provided error in the simulated R_g_ reflects the variability within the ensemble rather than a calculation error. **b**, Snapshots of the simulated trajectory. **c**, The P(r) for Pab1 at three salt concentrations are similar, consistent with the four charged RRM domains being largely non-interacting prior to condensation. **d**, The P(r) for the R123 construct (RRM1-RRM3 but lacking the flanking disordered regions) is nearly unchanged upon the addition of 0.5 M urea, further supporting a model where the RRMs already are largely non-interacting in the monomeric form without denaturant.

## Main References

1. Banani, S. F., Lee, H. O., Hyman, A. A. & Rosen, M. K. Biomolecular condensates: organizers of cellular biochemistry. Nat. Rev. Mol. Cell Biol. 18, 285–298 (2017).

2. Yoo, H., Triandafillou, C. & Drummond, D. A. Cellular sensing by phase separation: Using the process, not just the products. J. Biol. Chem. 294, 7151–7159 (2019).

3. Cherkasov, V. et al. Coordination of Translational Control and Protein Homeostasis during Severe Heat Stress. Current Biology 23, 2452–2462 (2013).

4. Jain, S. et al. ATPase-modulated stress granules contain a diverse proteome and substructure. Cell 164, 487–498 (2016).

5. Riback, J. A. et al. Stress-Triggered Phase Separation Is an Adaptive, Evolutionarily Tuned Response. Cell 168, 1028-1040.e19 (2017).

6. Kedersha, N. & Anderson, P. Stress granules: sites of mRNA triage that regulate mRNA stability and translatability. Biochem. Soc. Trans. 30, 963–969 (2002).

7. Wallace, E. W. J. et al. Reversible, Specific, Active Aggregates of Endogenous Proteins Assemble upon Heat Stress. Cell 162, 1286–1298 (2015).

8. Kato, M. et al. Cell-free Formation of RNA Granules: Low Complexity Sequence Domains Form Dynamic Fibers within Hydrogels. Cell 149, 753–767 (2012).

9. Molliex, A. et al. Phase Separation by Low Complexity Domains Promotes Stress Granule Assembly and Drives Pathological Fibrillization. Cell 163, 123–133 (2015).

10. Elbaum-Garfinkle, S. et al. The disordered P granule protein LAF-1 drives phase separation into droplets with tunable viscosity and dynamics. Proc. Natl. Acad. Sci. U.S.A. 112, 7189–7194 (2015).

11. Franzmann, T. M. & Alberti, S. Protein Phase Separation as a Stress Survival Strategy. Cold Spring Harb. Perspect. Biol. 11, a034058 (2019).

12. Jin, X. et al. Effects of pH alterations on stress- and aging-induced protein phase separation. Cell. Mol. Life Sci. 79, 380 (2022).

13. Kroschwald, S. et al. Different Material States of Pub1 Condensates Define Distinct Modes of Stress Adaptation and Recovery. Cell Rep. 23, 3327–3339 (2018).

14. Yoo, H., Bard, J. A. M., Pilipenko, E. V. & Drummond, D. A. Chaperones directly and efficiently disperse stress-triggered biomolecular condensates. Mol. Cell 82, 741-755.e11 (2022).

15. de Melo Neto, O. P., Standart, N. & de Sa, C. M. Autoregulation of poly(A)-binding protein synthesis in vitro. Nucleic Acids Res. 23, 2198–2205 (1995).

16. Melo, E. O., Dhalia, R., de Sa, C. M., Standart, N. & de Melo Neto, O. P. Identification of a C-terminal Poly(A)-binding Protein (PABP)-PABP Interaction Domain. J. Biol. Chem. 278, 46357–46368 (2003).

17. Cavanagh, J., Fairbrother, W. J., Palmer III, A. G. & Skelton, N. J. Protein NMR Spectroscopy: Principles and Practice. (Academic press, 1996).

18. Del Mar, C., Greenbaum, E. A., Mayne, L., Englander, S. W. & Woods, V. L. Structure and properties of α-synuclein and other amyloids determined at the amino acid level. Proc. Natl. Acad. Sci. U.S.A. 102, 15477–15482 (2005).

19. Englander, S. W., Mayne, L., Bai, Y. & Sosnick, T. R. Hydrogen exchange: the modern legacy of Linderstrøm-Lang. Protein Sci. 6, 1101–1109 (1997).

20. Boczek, E. E. et al. HspB8 prevents aberrant phase transitions of FUS by chaperoning its folded RNA-binding domain. eLife 10, e69377 (2021).

21. Lin, Y. et al. Redox-mediated regulation of an evolutionarily conserved cross-β structure formed by the TDP43 low complexity domain. Proc. Natl. Acad. Sci. U.S.A. 117, 28727–28734 (2020).

22. Englander, S. W., Sosnick, T. R., Englander, J. J. & Mayne, L. Mechanisms and uses of hydrogen exchange. Curr. Opin. Struct. Biol. 6, 18–23 (1996).

23. Zhang, Z. & Smith, D. L. Determination of amide hydrogen exchange by mass spectrometry: A new tool for protein structure elucidation: Amide hydrogen exchange by mass spectrometry. Protein Sci. 2, 522–531 (1993).

24. Mayne, L. et al. Many Overlapping Peptides for Protein Hydrogen Exchange Experiments by the Fragment Separation-Mass Spectrometry Method. J. Am. Soc. Mass Spectrom. 22, s13361-011-0235–4 (2011).

25. Bai, Y., Milne, J. S., Mayne, L. & Englander, S. W. Primary structure effects on peptide group hydrogen exchange. Proteins 17, 75–86 (1993).

26. Maris, C., Dominguez, C. & Allain, F. H.-T. The RNA recognition motif, a plastic RNA-binding platform to regulate post-transcriptional gene expression: The RRM domain, a plastic RNA-binding platform. FEBS J. 272, 2118–2131 (2005).

27. Hwang, T.-L., Mori, S., Shaka, A. J. & van Zijl, P. C. M. Application of Phase-Modulated CLEAN Chemical EXchange Spectroscopy (CLEANEX-PM) to Detect Water−Protein Proton Exchange and Intermolecular NOEs. J. Am. Chem. Soc. 119, 6203–6204 (1997).

28. Oxtoby, D. W. & Kashchiev, D. A general relation between the nucleation work and the size of the nucleus in multicomponent nucleation. J. Chem. Phys. 100, 7665–7671 (1994).

29. De Yoreo, J. J. & Vekilov, P. G. Principles of Crystal Nucleation and Growth. Rev. Mineral. Geochem. 54, 57–93 (2003).

30. Roder, H., Elöve, G. A. & Englander, S. W. Structural characterization of folding intermediates in cytochrome c by H-exchange labelling and proton NMR. Nature 335, 700–704 (1988).

31. Hoang, L., Bédard, S., Krishna, M. M., Lin, Y. & Englander, S. W. Cytochrome c folding pathway: kinetic native-state hydrogen exchange. Proc. Natl. Acad. Sci. U.S.A. 99, 12173–12178 (2002).

32. Lin, Y., Protter, D. S. W., Rosen, M. K. & Parker, R. Formation and Maturation of Phase-Separated Liquid Droplets by RNA-Binding Proteins. Mol. Cell 60, 208–219 (2015).

33. Harmon, T. S., Holehouse, A. S., Rosen, M. K. & Pappu, R. V. Intrinsically disordered linkers determine the interplay between phase separation and gelation in multivalent proteins. eLife 6, e30294 (2017).

34. Mittag, T. & Pappu, R. V. A conceptual framework for understanding phase separation and addressing open questions and challenges. Mol. Cell 82, 2201–2214 (2022).

35. Lindquist, S. The Heat-Shock Response. Annu. Rev. Biochem. 55, 1151–1191 (1986).

36. Morimoto, R. I. Proteotoxic stress and inducible chaperone networks in neurodegenerative disease and aging. Genes Dev. 22, 1427–1438 (2008).

37. Wang, S., Li, W., Liu, S. & Xu, J. RaptorX-Property: a web server for protein structure property prediction. Nucleic Acids Res. 44, W430–W435 (2016).

## References for methods section

38. Zmyslowski, A. M., Baxa, M. C., Gagnon, I. A. & Sosnick, T. R. HDX-MS performed on BtuB in E. coli outer membranes delineates the luminal domain’s allostery and unfolding upon B12 and TonB binding. Proc. Natl. Acad. Sci. U.S.A. 119, e2119436119 (2022).

39. Riback, J. A. et al. Innovative scattering analysis shows that hydrophobic disordered proteins are expanded in water. Science 358, 238–241 (2017).

40. Hopkins, J. B., Gillilan, R. E. & Skou, S. BioXTAS RAW: improvements to a free open-source program for small-angle X-ray scattering data reduction and analysis. J. Appl. Crystallogr. 50, 1545–1553 (2017).

41. Svergun, D. I. Determination of the regularization parameter in indirect-transform methods using perceptual criteria. J. Appl. Crystallogr. 25, 495–503 (1992).

42. Jumper, J. M., Faruk, N. F., Freed, K. F. & Sosnick, T. R. Accurate calculation of side chain packing and free energy with applications to protein molecular dynamics. PLoS Comput. Biol. 14, e1006342 (2018).

43. Jumper, J. M., Faruk, N. F., Freed, K. F. & Sosnick, T. R. Trajectory-based training enables protein simulations with accurate folding and Boltzmann ensembles in cpu-hours. PLoS Comput. Biol. 14, e1006578 (2018).

44. Peng, X. et al. Prediction and Validation of a Protein’s Free Energy Surface Using Hydrogen Exchange and (Importantly) Its Denaturant Dependence. J. Chem. Theory Comput. 18, 550–561 (2022).

## Reference for Extended Data figure legends

45. Kozlov, G. et al. Solution Structure of the Orphan PABC Domain from Saccharomyces cerevisiae Poly(A)-binding Protein. Journal of Biological Chemistry 277, 22822–22828 (2002).

46. Bremer, A. et al. Deciphering how naturally occurring sequence features impact the phase behaviours of disordered prion-like domains. Nat. Chem. 14, 196–207 (2022).

